# An unexpected alternative splicing of *SKU5-Similar3* in *Arabidopsis*

**DOI:** 10.1101/669507

**Authors:** Ke Zhou

## Abstract

Alternative splicing largely enhanced the diversity of transcriptome and proteome in eukaryas. Along with technical development, more and more alternatively splicing was demonstrated. Here, we report an unexpected alternative splicing of *SKU5-Similar 3* (*SKS3*) within a special splicing site in *Arabidopsis*. Based on bioinformatics database, *SKS3* was predicted to be alternatively transcribed into two variants, *SKS3.1* and *SKS3.2*, which encoded a GPI-anchored protein and a soluble secretory protein respectively. But, instead of *SKS3.2*, a novel variant, *SKS3.3*, which encoded a protein with transmembrane region at its C-terminus, was demonstrated based on our experimental data. Interestingly, it exhibited a different organ-specific expression pattern from *SKS3.1*, and its intron splicing site did not follow ‘GT-AG’ rule or any reported rules.

## Introduction

In eukaryas, precursor messenger RNA (pre-mRNA) was transcribed directly from genes, and then spliced under a conserved mechanism, where exons were spliced to be mature mRNA for translation, and introns were removed ^1–3^. The splicing sites were not always unique, but could be altered, which resulted in the facts that, one gene could be transcribed into several variants, and then encoded several proteins ^4^. It largely enhanced the diversity of transcriptome and proteome ^5,6^. Along with the technological development, more and more alternative splicing was identified. Based on RNA-seq data, >95– 100% of human genes ^3^, and up to 60% of intron-containing genes of *Arabidopsis* genes ^1,2,5,7^, are alternatively spliced and could generate at least two alternative variants. The alternative splicing possesses were demonstrated to be performed by the protein-RNA complexes, spliceosomes, which could recognize the 5’, 3’ splice sites and the branch point of introns ^8–10^. This special recognition and splicing resulted in the “GT-AG” rule of introns, which means the spliced introns mostly start from “GT” and end at “AG” ^4,8^.

Interestingly, the alternative splicing of genes was not consistent, but its occurrence and the expression pattern of alternatively spliced variants were reported to be regulated by various abiotic stresses ^11–15^, at developmental stages^16^, or in different organs^17^. Although its exact mechanism hasn’t been well revealed yet ^18^, a few reports indicated its importance during development, such as, the different alternative splicing variants of HAB1 play opposite roles during ABA signaling in *Arabidopsis* ^19^.

*SKU5-Similar 3* (*SKS3*) encodes a plasma membrane attached glycosylphosphatidylinositol-anchored protein, which belongs to a SKU5-Similar protein subfamily that were redundantly essential for cell polar expansion and cell wall synthesis of roots in *Arabidopsis* ^20^. In this study, we reported an unexpected organ-specific alternatively spliced variant of *SKS3* in *Arabidopsis*, which could encode a plasma membrane attached protein with transmembrane region at its C-terminus. Interestingly, its splicing site was unique, which did not follow the “GT-AG” rule, or any other reported rules. But due to the short repeat “ATCCATC” localized close to the border of spliced intron, the exact site could not be identified. The reverse gene at its 3’-terminal, which could encode a long noncoding RNA (lncRNA), might participate in these processes.

## METIRIALS AND METHODS

### RNA extraction and semi-quantitative RT-PCR

RNA was extract from 5-day old seedlings, rossetta leaves, whole opening flowers and mature siliques from *Arabidopsis*, cDNA was synthesized by Oligo d(T).

### Primers

Primers utilized for semi-quantitative PCR: SKS3-F, TTTTCTCCATTTTCACTCACTGCT; SKS3-R, CTAATATGATATCCGATCCCGGTT; Actin-F, GTTAGCAACTGGGATGATATGG; Actin-R, CAGCACCAATCGTGATGACTTGCCC.

## RESULTS

### Predicted alternative splicing of *SKS3* gene

According to NCBI (https://www.ncbi.nlm.nih.gov/) and TAIR (http://www.arabidopsis.org/) database, two transcriptional variants, *SKS3.1 and SKS3.2* were predicated to be transcribed from *SKS3* gene. *SKS3.1* encodes a protein precursor containing 589 amino acid residues, which could be modified with a glycosylphosphatidylinositol modification at C-terminus; *SKS3.2* encodes a protein precursor containing 614 amino acid residues, which was predicted soluble and to enter the secretory pathway. Gene structures of *SKS3.1* and *SKS3.2* showed that, the predicted alternative splicing occurred at the last exon where the STOP codon was lost in *SKS3.2* (Fig.1).

**Fig 1.**
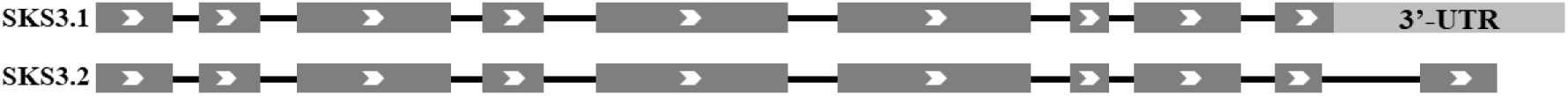
Gene structures of *SKS3.1* and *SKS3.2* predicted by NCBI and TAIR database. Arrows showed the translation direction; exons were in dark grey; untraslated region (UTR) was in light grey; and introns were shown as black lines.

### Observed alternative splicing of *SKS3* gene *in vivo*

To precise the alternative splicing of *SKS3* in *Arabidopsis, SKS3* variants were cloned from cDNA generated from different organs. Surprisingly, instead of *SKS3.2* variant, a novel variant, *SKS3.3*, was identified to exhibit a different expression pattern with *SKS3.1* (Fig.2A). Through analyzing the gene structure of *SKS3.3*, an unusual alternative splicing and a long intron, which include the whole 6^th^, 7^th^ and 8^th^ exons, partly 5^th^ and 9^th^ exons, and all introns between them, were revealed (Fig.2C). Interestingly, the variant *SKS3.3* encodes a shorter precursor containing 314 amino acid residues, which was predicted to be a transmembrane protein with a short transmembrane region at its C-terminus (Fig.2B).

**Fig 2.**
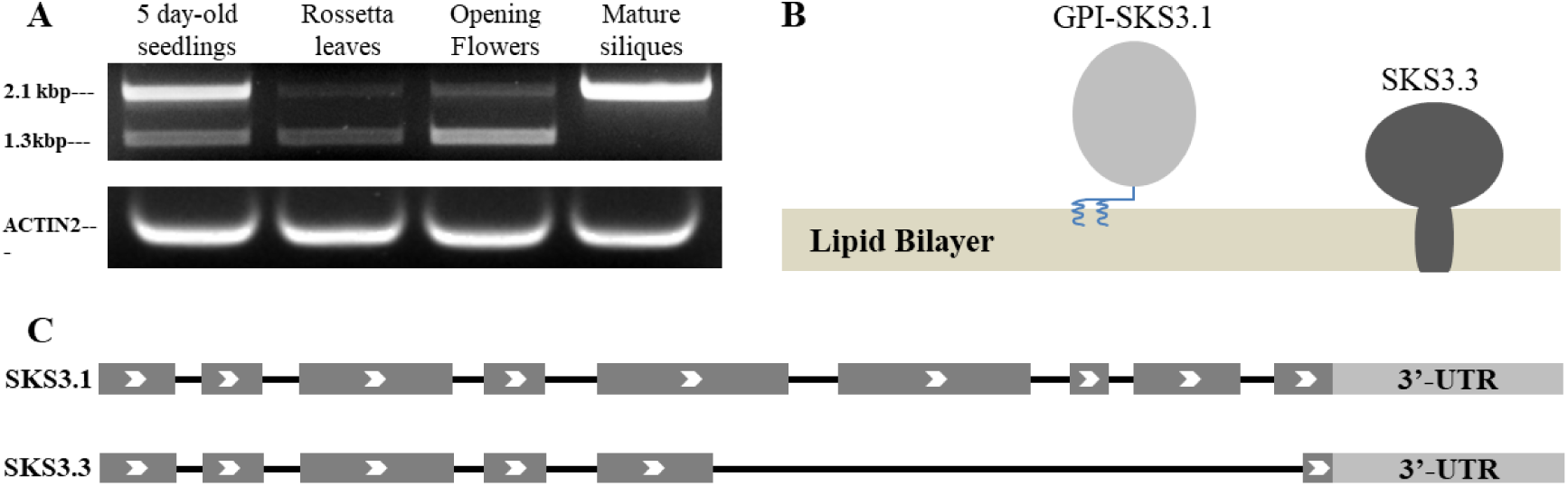
Alternatively spliceing of *SKS3* gene in *Arabidopsis*. **A**, two *SKS3* variants were amplified from cDNA extracted from different organs of *Arabidopsis*; Actin was utilized as reference and primers were indicated in methods; **B**, predicted GPI-anchored protein SKS3.1 and transmembrane protein SKS3.3; **C**. gene structure of *SKS3.1* and *SKS3.3*. Arrows showed the translation direction; exons were in dark grey; untraslated region (UTR) was in light grey, and introns were shown as black lines.

### The novel splicing site within “ATCCATC” of *SKS3.3*

Interestingly, instead of following “GT-AG” rule, the borders of the spliced intron did not follow any reported splicing rule, but with an unusual “ATCCATC” repeat close to the splicing site (Fig. 3). But due to the presence of this repeat, the exact splicing site of *SKS3. 3* could not be recognized, but could only be limited within the short repeat “ATCCATC”.

**Fig.3.**
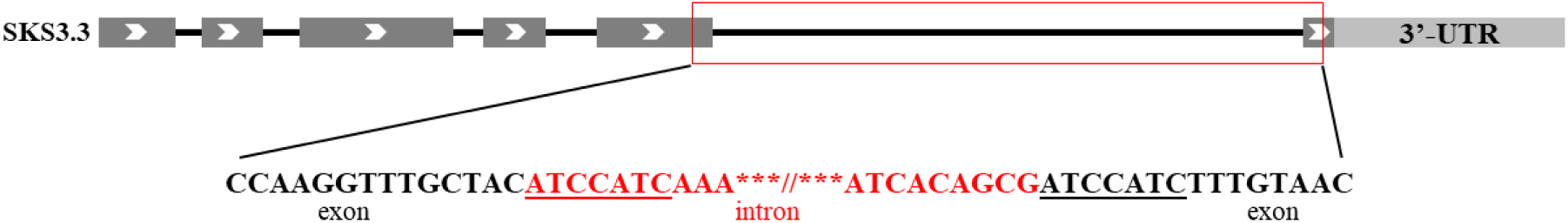
Splicing site of *SKS3.3* variant. Arrows showed the translation direction; exons were in dark grey; untraslated region (UTR) was in light grey, and introns were shown as black lines, the alternatively spliced intron was in red blank and the sequence close to splicing site was labeled in red, and the “ATCCATC” repeat was underlined.

## DISCUSSION

Alternative splicing largely enhanced the diversity of transcriptome and proteome, which allows one gene to encode more proteins. According to NCBI and TAIR database, *SKS3* gene was predicted to be transcribed into two variants, *SKS3.1* and *SKS3.2*, which encode a GPI-anchored protein and a soluble protein respectively. But according to our experimental data, two variants, *SKS3.1* and *SKS3.3*, which exhibited different expression patterns in various organs, were identified, but not *SKS3.2*. It indicated the complexity of alternative splicing in *Arabidopsis*.

In our study, the two identified variants of *SKS3* were predicted to encode a GPI-anchored protein and a smaller transmembrane protein, which were both attached to the outer surface of plasma membrane eventually. It reminded us of the two variants of HAB1, which could encode two proteins act opposite roles during ABA signaling pathway in *Arabidopsis*^19^, and suggested a potential functional diversity of SKS3.1 and SKS3.3 protein.

Through investigating the gene structure of *SKS3*, a reverse gene *AT5G01745* that encode a lncRNA was found at its 3’-terminus, overlapping with the alternative splicing site of *SKS3.3* variant (Fig.4). Long noncoding RNA (lncRNA), has been reported to be involved in alternative splicing, potentially through forming double-strand to prevent the recognition and splicing from spliceosome ^21–25^. It suggested the involvement of lncRNA in the unexpected alternative splicing, and it would be very interesting to further investigate the connection between the alternative splicing and the lncRNA, and the regulation of this lncRNA.

**Fig.4.**
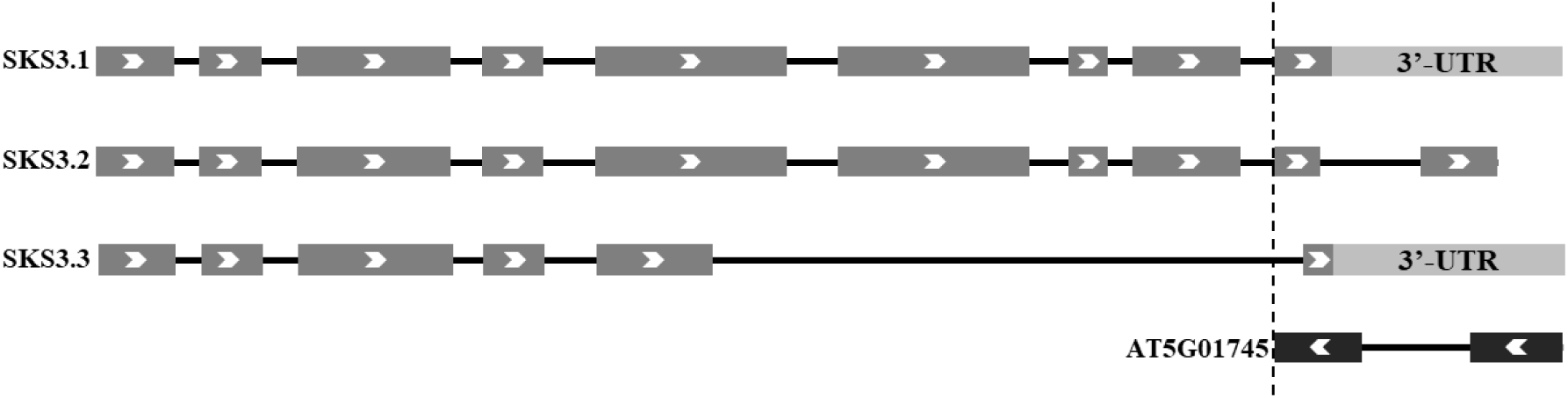
Transcriptional variants of *SKS3*, and the lncRNA encoding gene *AT5G01745* reversely localized at 3’-terminus of *SKS3*. Arrows showed the translation direction; exons were in dark grey; untraslated region (UTR) was in light grey, and introns were shown as black lines.

Generally, due to the recognition and splicing from spliceosomes, the vast majority of intron splicing in *Arabidopsis* followed the “GT-AG” rule, which means the spliced introns mostly start from “GT” and end at “AG” ^4,8–10^, with a few exceptions, such as “GC-AG” rule ^5^. Interestingly, an unusual intron border was identified in *SKS3.3* which has not been reported. But due to the presence of the repeated “ATCCATC” close to the splicing site of the unexpected intron, the exact splicing site of *SKS3.3* could not be recognized, but could only be limited within the short repeat “ATCCATC”. It would be not surprisingly if we found the involvement of lncRNA in its alternative splicing through forming double strand RNA with 3’-terminus of *SKS3* pre-mRNA.

## Acknowledgments

Ke ZHOU designed and performed the experiments, wrote and revised the manuscript.

## References

1. Bush, S. J., Chen, L., Tovar-Corona, J. M. & Urrutia, A. O. Alternative splicing and the evolution of phenotypic novelty. Philos. Trans. R. Soc. B Biol. Sci. 372, 1–7 (2017).

2. Reddy, A. S. N., Marquez, Y., Kalyna, M. & Barta, A. Complexity of the Alternative Splicing Landscape in Plants. Plant Cell 25, 3657–3683 (2013).

3. Lee, Y., Rio, D. C., Biology, S. & Biology, C. Mechanisms and Regulation of Alternative PremRNA Splicing. Annu.Rev.Biochem. 291–323 (2015). doi:10.1146/annurev-biochem-060614-034316.Mechanisms

4. Kelemen, O. et al. Function of alternative splicing. Gene 514, 1–30 (2013).

5. Filichkin, S. A. et al. Genome-wide mapping of alternative splicing in Arabidopsis thaliana. Genome 45–58 (2010). doi:10.1101/gr.093302.109.2008

6. Severing, E. I., van Dijk, A. D. J. & Van Ham, R. C. H. J. Assessing the contribution of alternative splicing to proteome diversity in Arabidopsis thaliana using proteomics data. BMC Plant Biol. 11, 82 (2011).

7. Wang, Z., Gerstein, M. & Snyder, M. RNA-Seq: a revolutionary tool for transcriptomics. Nat Rev Genet. 10, 57–63 (2009).

8. Gooding, C. et al. A class of human exons with predicted distant branch points revealed by analysis of AG dinucleotide exclusion zones. Genome Biol. 7, 1–19 (2006).

9. Behzadnia, N. et al. Composition and three-dimensional EM structure of double affinity-purified, human prespliceosomal A complexes. EMBO J. 26, 1737–1748 (2007).

10. Dai, Q. et al. RNA catalyses nuclear pre-mRNA splicing. Nature 503, 229–234 (2013).

11. Zhu, G., Li, W., Zhang, F. & Guo, W. RNA-seq analysis reveals alternative splicing under salt stress in cotton, Gossypium davidsonii. BMC Genomics 19, 1–15 (2018).

12. Leviatan, N., Alkan, N., Leshkowitz, D. & Fluhr, R. Genome-Wide Survey of Cold Stress Regulated Alternative Splicing in Arabidopsis thaliana with Tiling Microarray. PLoS One 8, (2013).

13. Li, W., Lin, W.-D., Ray, P., Lan, P. & Schmidt, W. Genome-Wide Detection of Condition-Sensitive Alternative Splicing in Arabidopsis Roots. Plant Physiol. 162, 1750–1763 (2013).

14. Yeoh, L. M. et al. A serine-arginine-rich (SR) splicing factor modulates alternative splicing of over a thousand genes in Toxoplasma gondii. Nucleic Acids Res. 43, 4661–4675 (2015).

15. Kwon, Y. J., Park, M. J., Kim, S. G., Baldwin, I. T. & Park, C. M. Alternative splicing and nonsense-mediated decay of circadian clock genes under environmental stress conditions in Arabidopsis. BMC Plant Biol. 14, 1–15 (2014).

16. Deschênes, M. & Chabot, B. The emerging role of alternative splicing in senescence and aging. Aging Cell 16, 918–933 (2017).

17. Loraine, A. E., McCormick, S., Estrada, A., Patel, K. & Qin, P. RNA-Seq of Arabidopsis Pollen Uncovers Novel Transcription and Alternative Splicing. Plant Physiol. 162, 1092–1109 (2013).

18. Cui, Z. et al. SKIP controls flowering time via the alternative splicing of SEF pre-mRNA in Arabidopsis. BMC Biol. 15, 1–17 (2017).

19. Wang, Z. et al. ABA signalling is fine-tuned by antagonistic HAB1 variants. Nat. Commun. 6, 8138 (2015).

20. Zhou, K. GPI-anchored SKS proteins regulate root development through controlling cell polar expansion and cell wall synthesis. Biochem. Biophys. Res. Commun. 509, (2019).

21. Chekanova, J. A. Long non-coding RNAs and their functions in plants. Curr. Opin. Plant Biol. 27, 207–216 (2015).

22. Bardou, F. et al. Long Noncoding RNA Modulates Alternative Splicing Regulators in Arabidopsis. Dev. Cell 30, 166–176 (2014).

23. Severing, E. et al. Arabidopsis thaliana ambient temperature responsive lncRNAs. BMC Plant Biol. 18, 1–10 (2018).

24. Lu, Z. et al. Identification and characterization of novel lncRNAs in Arabidopsis thaliana. Biochem. Biophys. Res. Commun. 488, 348–354 (2017).

25. Ma, L., Bajic, V. B. & Zhang, Z. On the classification of long non-coding RNAs. RNA Biol. 10, 924–933 (2013).

